# Dynamic Changes in Chloride Homeostasis Coordinate Midbrain Inhibitory Network Activity during Reward Learning

**DOI:** 10.1101/2024.11.18.624156

**Authors:** Joyce Woo, Ajay Uprety, Daniel Reid, Irene Chang, Aelon Ketema Samuel, Helena de Carvalho Schuch, Caroline C Swain, Alexey Ostroumov

## Abstract

The ability to associate environmental stimuli with positive outcomes is a fundamental form of learning. While extensive research has focused on the response profiles of midbrain dopamine neurons during associative learning, less is known about learning-mediated changes in the afferents that shape their responses. We demonstrate that during critical phases of learning, anion homeostasis in midbrain GABA neurons – a primary source of input to dopamine neurons – is disrupted due to downregulation of the chloride transporter KCC2. This alteration in GABA neurons preferentially impacted lateral mesoaccumbal dopamine pathways and was not observed after learning was established. At the network level, learning-mediated KCC2 downregulation was associated with enhanced synchronization between individual GABA neurons and increased dopamine responses to reward-related stimuli. Conversely, enhancing KCC2 function during learning reduced GABA synchronization, diminished relevant dopamine signaling, and prevented cue-reward associations. Thus, circuit-specific adaptations in midbrain GABA neurons are crucial for forming new reward-related behaviors.

## Introduction

An organism’s survival and adaptation depend on its ability to learn associations between environmental stimuli and rewards. Associative reward learning links events to outcomes, and is a fundamental process essential to adaptive behaviors; its disruption is a core feature of many debilitating neuropsychiatric disorders, such as major depressive disorder^1^, addiction^2,3^, and schizophrenia^4^.

Seminal work has shown that the rate of associative reward learning relies on phasic dopamine (DA) signaling within mesolimbic pathways projecting from the ventral tegmental area (VTA) targeting the nucleus accumbens (NAc)^5,6^. These learning-related DA responses exhibit remarkably focused circuit-specificity, in which reward and cue-related phasic activation dominate in lateral VTA DA neurons that project to the lateral subregions of the NAc^5,7–9^. During initial phases of learning, these neurons exhibit transient firing primarily in response to reward presentations. However, following repeated cue-reward pairings, DA neurons begin to exhibit additional bursts of activity at cue onset, indicating the strength of association between rewards and environmental stimuli^6,10,11^. Despite extensive research involving DA and reward encoding in the brain, the mechanisms shaping these representations are still unclear.

GABA-releasing neurons in the VTA are key regulators of VTA DA neurons^12,13^. These VTA GABA neurons increase firing during cues that predict rewards^13–15^, and exhibit drastic alterations in transmission after exposure to drugs of abuse^16–19^. However, there has been surprisingly little focus on how enhanced GABAergic signaling contributes to the acquisition of reward-associated behaviors.

Recent studies have shown that highly salient experiences, such as stress, addictive drugs, and chronic pain, induce a shift toward excitatory GABA_A_ receptor signaling in VTA GABA neurons^18–20^. This shift in GABA signaling stems from a downregulation in the neuron-specific Cl^−^ extrusion pump, KCC2, which primarily functions to maintain low intracellular Cl^−^ levels. While midbrain DA neurons exhibit little to no expression of KCC2^21^, VTA GABA neurons are significantly affected by KCC2 impairment. The resulting intracellular Cl^−^ accumulation leads to depolarizing shifts in the Cl^−^ reversal potential (E_GABA_), and compromises GABA_A_ receptor-mediated inhibition^20,22^. Ultimately, functional KCC2 downregulation in VTA GABA neurons following stress and drug exposure enhances excitatory GABA_A_ receptor signaling and amplifies the acquisition of drug-taking behaviors^17,19,20^. Changes in chloride homeostasis within the VTA circuitry may reflect a pathological usurpation of mechanisms that normally support reward-related learning – a framework that has previously been proposed for addiction^23^. Despite its previous implication during pathological states, intracellular chloride regulation during naturalistic learning contexts has not yet been examined.

To address this gap, we show that dynamic changes in intracellular anion homeostasis in GABAergic signaling emerge during reward learning and thereby shape the activity of downstream VTA DA targets. Utilizing behavioral, electrophysiological, and molecular approaches, we demonstrate that VTA GABA neurons exhibit altered Cl^−^ homeostasis through functional KCC2 downregulation within fine temporal windows during reward learning. Importantly, we show that enhancing KCC2 function in the VTA halts the progression of cue-reward learning. Moreover, our findings indicate that functional KCC2 downregulation is pathway-specific and plays a crucial role in synchronizing the activity of midbrain GABAergic networks. Finally, we show that synchronized GABAergic activity can amplify phasic firing in DA neurons. These findings indicate that circuit-specific alterations in KCC2 are a fundamental mechanism that sculpts experience-induced circuit remodeling and reward learning.

## Results

### Learning-dependent downregulation of KCC2 in VTA GABA neurons

We first investigated whether reward learning altered anion homeostasis in VTA GABA neurons. Given previous findings indicating that depolarizing shifts in E_GABA_ are calcium (Ca^2+^)-dependent^24^, we focused selectively on GABAergic populations that were activated during cue-reward pairings. To achieve this, male and female GAD-Cre rats received intra-VTA injections of Cre-dependent blue fluorescent protein (BFP) and a fluorescent Ca^2+^ sensor CaMPARI2 that converts from green to red fluorescence in the presence of high calcium concentrations and 405 nm (UV) light^25^ (Fig. 1A and Extended Data Fig. 1A). In freely moving rats undergoing Pavlovian conditioning (see Methods), the UV illumination was delivered through an implanted optic fiber during the 5-second auditory tone (conditioned stimulus, CS) and subsequent sugar pellet presentations (unconditioned stimulus, US, Fig. 1B, Paired Conditioning). In a separate unpaired conditioning group, the UV light was also delivered throughout the CS and US, but these two stimuli were never presented in immediate succession (Fig. 1B, Unpaired Conditioning). Compared to the unpaired group, reward-seeking behaviors during CS presentations significantly increased after CS-US pairing over the 13-day period (Fig. 1C), indicating the formation of cue-reward association. Sex differences were not detected in the paired group (Extended Data Fig. 1B).

**Figure 1.**
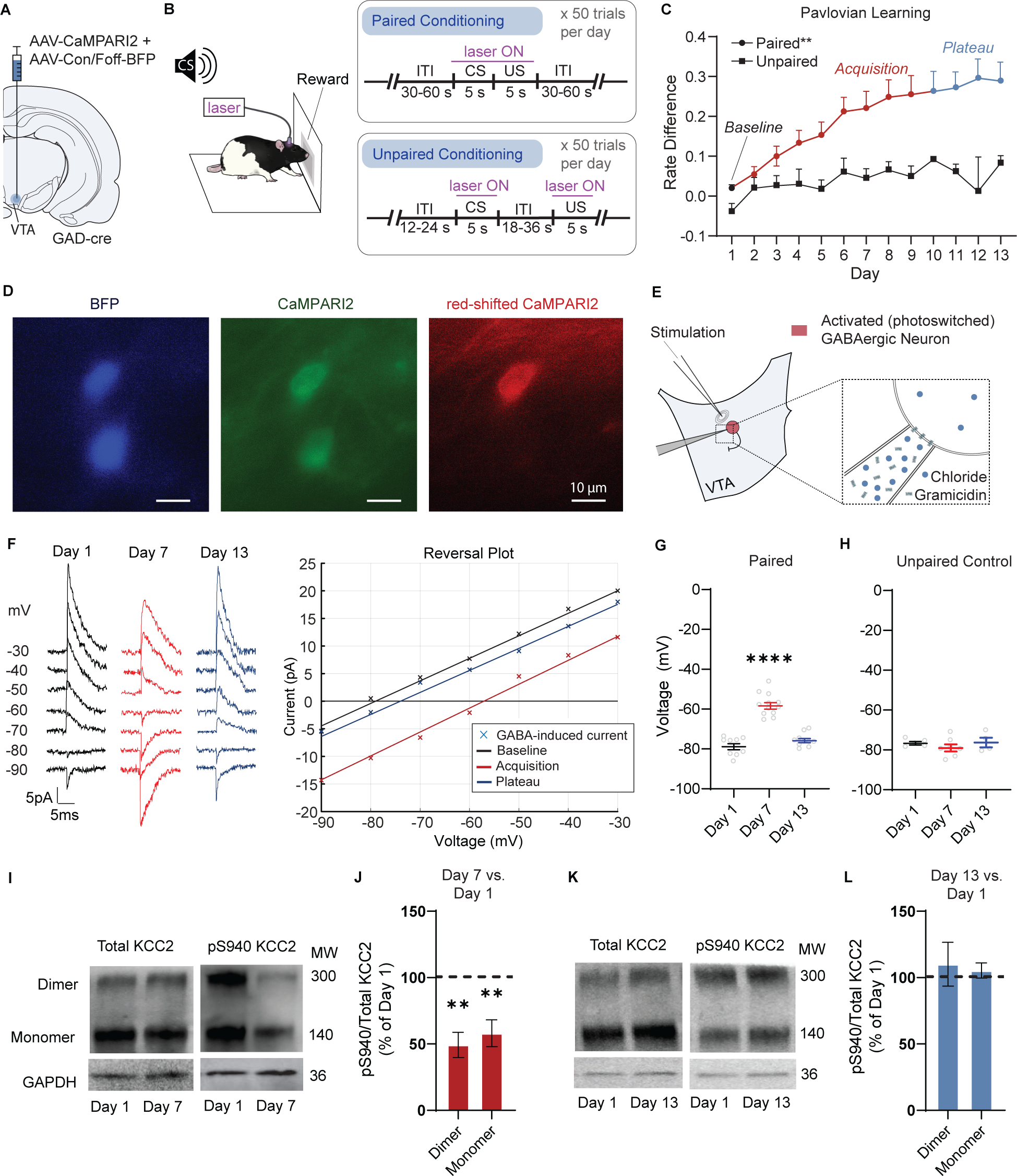
Learning-dependent downregulation of KCC2 in VTA GABA neurons. A) Cre-dependent BFP and CaMPARI2 were injected in the VTA of GAD-Cre rats. B) During the paired Pavlovian learning task, animals were exposed to 50 conditioning trials consisting of a 5-second auditory tone (conditioned stimulus, CS), followed by delivery of a sugar pellet (unconditioned stimulus, US), and an intertrial-interval ranging from 30-60 seconds. To identify VTA GABA cells that were activated during cue-reward association, laser UV illumination was applied during CS and US. An unpaired training was introduced as a control procedure for associative learning wherein CS and US were presented in the same trial but separated in time. In the unpaired group, UV stimulation was applied during both cue and reward. C) In contrast to unpaired control, the paired group showed progression of reward learning quantified as a change in number of port entries during CS relative to 5-seconds preceding CS (i.e., rate difference). The conditioning time course in the paired group consisted of baseline, acquisition, and plateau phases. **p < 0.01, mixed-effect model, group x time, F(12, 395) = 2.336, n = 13-39 rats. D) BFP-expressing VTA GABA neurons expressing green and red fluorescence after 7 days of Pavlovian training. E) E_GABA_ in photoconverted VTA GABA neurons was recorded using gramicidin perforated patch clamp. GABA_A_ eIPSCs were evoked by electrical stimulation in the presence of ionotropic glutamate and GABA_B_ receptor antagonists. F) Left: representative eIPSCs recording from baseline (black), acquisition (red), and plateau (blue) periods of Pavlovian conditioning at the given holding potentials. For display, the traces were filtered, and stimulus artifacts were removed. Right: E_GABA_ was determined from an I-V curve as a membrane voltage at which eIPSCs exhibited zero amplitude. G) VTA GABA neurons from paired groups demonstrated significantly more depolarized E_GABA_ during acquisition periods compared to baseline and plateau: −58.08 ± 1.64 mV on days 5-7 (red) versus −78.89 ± 1.52 mV on day 1 (black), and −75.84 ± 1.04 mV on day 13 (blue). ****p < 0.0001, one-way ANOVA, F = 62.39, n = 9-12 cells, 5-7 rats. H) In unpaired groups, E_GABA_ did not show significant change across learning phases (p = 0.49, one-way ANOVA, n = 4-7 cells, 2-3 rats). I) Western blot analysis was conducted for total KCC2 and phosphorylated KCC2 (pS940) with GAPDH as a loading control. A representative western blot indicates no differences in total KCC2 expression after 7 days of learning. However, during acquisition, animals showed reduced expression of pS940 KCC2 relative to total KCC2 when compared to day 1. J) Densitometric analysis of western blot showed significantly reduced expression of pS940 KCC2 for animals in acquisition periods of learning compared to baseline: 49.31 ± 9.46% for dimers, 58.03 ± 10.13% for monomers. **p < 0.01, paired t-test, n = 6. K) A representative western blot indicating no differences in either total or phosphorylated KCC2 expression after 13 days of learning compared to day 1. L) Expression of pS940 KCC2 for animals in plateau periods shows no significant difference from baseline (p = 0.82 for dimers, p = 0.30 for monomers, n = 8 rats, paired t-test).

Upon examination of the paired conditioning time course, we delineated three distinct phases of learning: *baseline*—referring to the first day of learning, *acquisition*—characterized by a rapid ascent in rate difference, and *plateau* – a period of stable responding with <10% difference in performance over 3 consecutive days (Fig. 1C). To examine whether these phases impacted anion homeostasis in VTA GABA neurons, we measured the reversal potential for GABA_A_ receptor mediated currents (E_GABA_) in cells expressing BFP and photoconverted red CaMPARI (Fig. 1D). After days 1, 5-7, and 13 of Pavlovian conditioning, we performed gramicidin-perforated patch-clamp recordings to preserve the intracellular anion concentrations (Fig. 1E). E_GABA_ was determined as the membrane potential at which evoked inhibitory postsynaptic currents (eIPSCs) change their direction from inward to outward (Fig. 1F). During the acquisition phase of Pavlovian conditioning, VTA GABA neurons in male and female rats showed significantly more depolarized E_GABA_ values compared to baseline and plateau phases of learning (Fig. 1G and Extended Data Fig. 1C). Although E_GABA_ was depolarized during acquisition, Pavlovian learning did not alter the membrane resting potential in VTA GABA neurons (Extended Data Fig. 1D). In the unpaired conditioning group, red-shifted CaMPARI expressing VTA GABA neurons maintained hyperpolarized E_GABA_ values across all timepoints (Fig. 1H).

A depolarizing shift in E_GABA_ reflects reductions in Cl^−^ extrusion capacity, which has been previously associated with dephosphorylation of KCC2 at serine 940 (S940)^20,26,27^. To examine whether dephosphorylation of KCC2 was altered during reward learning, we performed western blot analysis using antibodies against total KCC2 and phospho-S940 KCC2 at distinct phases of Pavlovian conditioning. Our immunoblots revealed prominent bands at 140 and 270 kDa for both total and phospho-S940 KCC2, indicating the presence of monomeric and dimeric forms of KCC2 (Fig. 1I). As anticipated, the ratio of phosphorylated S940 KCC2 to total KCC2 during acquisition of learning (days 5-7) was significantly lower compared to baseline (day 1, Fig. 1J). In contrast, no significant differences in the ratio of phosphorylated S940 KCC2 to total KCC2 were observed between baseline and plateau groups (day 13, Figs. 1K and 1L). We observed no significant differences in the expression of phospho-S940 across sexes (Extended Data Fig. 1E), and in the expression of total KCC2 protein between different phases of learning (Extended Data Fig. 1F). Given the previous immunolabeling studies showing that KCC2 protein is expressed in non-DA, GABAergic neurons within the VTA^19,21^, our results suggest that dephosphorylation of KCC2 protein at S940 decreases KCC2 function in VTA GABA neurons only during the acquisition of cue-reward association.

### Learning-mediated KCC2 downregulation impacts NAc lateral shell projecting DA neurons

Next, we examined whether depolarizing shifts in E_GABA_ within VTA GABA neurons altered GABA release on distinct DA neuron projections to different subregions of the NAc. To label projection-specific DA neurons, we injected a retrogradely transported Cre-dependent AAV vector containing GFP in the NAc lateral shell, medial shell, and core of male and female TH-Cre rats (Fig. 2A)^5,28,29^. Then, we assessed learning-related changes in GABA release, by measuring the frequency of spontaneous inhibitory postsynaptic currents (sIPSCs) in GFP-expressing VTA DA neurons immediately after days 1, 5-7, and 13 of Pavlovian conditioning (Fig. 2B).

**Figure 2.**
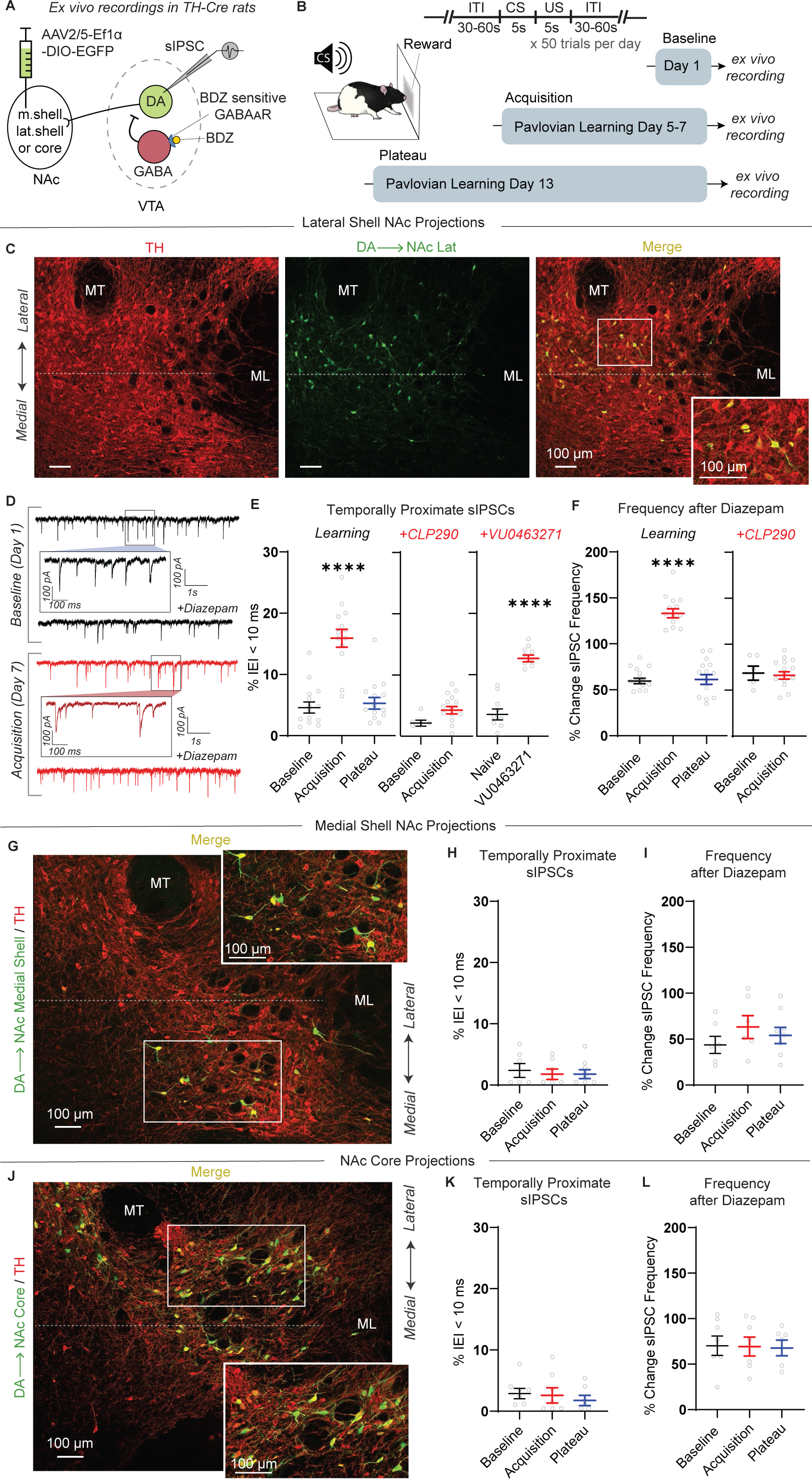
Learning-mediated KCC2 downregulation impacts NAc lateral shell projecting DA neurons. A) AAV expressing a Cre-dependent eGFP was injected in either NAc lateral, medial shell, or core of TH-Cre rats to retrogradely label DA neuron projections. SIPSCs onto eGFP-expressing VTA DA neurons were recorded using the whole-cell patch-clamp configuration. Diazepam was applied to potentiate GABA_A_ receptor function in VTA GABA neurons. B) After AAV injections, animals underwent paired Pavlovian conditioning, and horizontal slices were collected for *ex vivo* electrophysiology at different stages of learning. C) Immunostaining showing that Cre-dependent eGFP injection in the NAc lateral shell of TH-Cre rats led to eGFP (green) expression in lateral VTA DA cells (TH, red). D) Representative recordings of sIPSCs before and after diazepam application after day 1 (black) and day 7 (red) of learning. Insets demonstrate learning-mediated changes in the distribution of sIPSC inter-event intervals. E) Left: Lateral shell-projecting DA neurons demonstrate a significant increase in sIPSCs with interevent intervals of <10 ms during acquisition (15.90 ± 1.46% of all sIPCSs, red) compared to baseline (4.59 ± 0.91%, black) and plateau (5.28 ± 0.95%, blue) phases of learning. ****p < 0.0001, one-way ANOVA, F= 31.39, n = 13-14 cells, 3-4 rats. Center: Following CLP290 incubation, the number of sIPSCs with interevent intervals of <10 ms did not differ between baseline (2.16 ± 0.49%, black) and acquisition of learning (4.26 ± 0.6%, red, p = 0.06, t-test, n = 5-14 cells, 2-4 rats). Right: Bath application of VU0462371 on VTA slices from naïve animals increased the percentage of sIPSCs with interevent intervals of <10 ms in lateral shell-projecting DA neurons up to acquisition levels: 12.67 ± 0.55% (compared to pre-VU0462371: 3.53 ± 0.89%). ****p < 0.0001, t-test, n = 9 cells, 3 rats. F) After acquisition of learning (red), lateral shell-projecting DA neurons showed significantly increased diazepam-mediated sIPSC frequencies (133.2 ± 4.87%) compared to neurons from baseline (59.58 ± 3.07%) and plateau (61.23 ± 5.27%). ****p < 0.0001, one-way ANOVA, F = 85.37, n = 13-14 cells, 3-4 rats. Following CLP290 incubation, diazepam-induced sIPSCs frequency did not differ between baseline (73.36 ± 8.21%, black) and acquisition of learning (66.08 ± 4.06%, red, p = 0.39, t-test, n = 5-14 cells, 2-4 rats). G) Immunostaining from the VTA showing TH (red) and eGFP (green) labeling following Cre-dependent eGFP injection in the NAc medial shell of TH-Cre rats. H) Quantified percentages of sIPSCs with interevent intervals of < 10 ms in NAc medial shell-projecting DA neurons revealed no differences across learning (p = 0.86, one-way ANOVA, n = 6-8 cells, 4 rats. I) NAc medial shell-projecting DA neurons showed no significant differences in diazepam-induced sIPSC frequency across learning (p = 0.45, one-way ANOVA, n = 6-8 cells, 4 rats. J) TH (red) and eGFP (green) immunolabeling in the VTA of TH-Cre rats after Cre-dependent eGFP expression in the NAc core. K) No learning-mediated differences in the percentage of sIPSCs with interevent intervals of <10 ms were noted among NAc core-projecting DA neurons at different phases of learning. (p = 0.73, one-way ANOVA, n = 6-7 cells, 3-5 rats). L) Following CLP290 incubation, diazepam-induced sIPSC frequency in NAc core-projecting DA neurons did not differ between baseline and acquisition of learning (p = 0.64, n = 6-7 cells, 3-5 rats, one-way ANOVA).

Lateral shell-projecting DA neurons were located predominantly in the lateral portions of the VTA (Fig. 2C). While sIPSC frequency did not change across learning (Extended Data Fig. 2A), we observed an increase in spontaneous GABA release events occurring in close temporal proximity to each other during acquisition (Fig. 2D, top black and red traces with insets). Quantitative analysis revealed that after days 5-7 of Pavlovian conditioning, lateral shell-projecting DA neurons exhibited a significantly higher percentage of sIPSCs with interevent intervals < 10 ms, compared to days 1 and 13 (Fig. 2E, left). To examine the role of KCC2 downregulation in this effect, we recorded sIPSCs in slices incubated with pharmacological KCC2 activator CLP290 (∼1 h, 10 µM)^18,19^. CLP290 treatment after days 5-7 of learning decreased the percentage of sIPSCs with interevent intervals of < 10 ms to the values observed at days 1 and 13 (Fig. 2E, middle). Furthermore, lateral shell-projecting DA neurons from naïve control rats treated with KCC2 inhibitor VU0463271 (10 µM) showed a potentiation of closely occurring sIPSCs that was similar to days 5-7 of learning (Fig. 2E, right). Importantly, neither CLP290 nor VU0463271 altered the total number of sIPSCs per second (Extended Data Figs. 2B and 2C). Increased number of sIPSCs occurring in close temporal proximity, without changes in the total number of events, suggests enhanced synchronized GABA release on a single DA neuron^30,31^.

To confirm the impact of KCC2 modulation on GABA release onto DA neurons, we next used a pharmacological approach. Prior studies showed that KCC2 downregulation leads to excitation of VTA GABA neurons during intense GABA_A_ receptor stimulation^14^. Compared to DA neurons, GABA_A_ receptors in VTA GABA neurons are much more sensitive to benzodiazepines due to differences in subunit composition^32^. Thus, benzodiazepines, such as diazepam, potentiate GABA_A_ receptor function primarily in VTA GABA, but not in DA neurons. Indeed, diazepam was previously shown to suppress VTA GABA neuron activity and to attenuate GABA release onto DA neurons^17,33^. Upon KCC2 downregulation, however, diazepam-mediated potentiation of GABA_A_ receptor signaling excited VTA GABA neurons and enhanced GABA release onto DA neurons^17^. Consistent with this model, bath-application of diazepam (5 µM) during the acquisition phase of learning significantly increased sIPSC frequency in DA neurons compared to baseline and plateau (Figs. 2D and 2F, left). Incubation with CLP290 prevented diazepam-mediated increase in sIPSC frequency (Fig. 2F, right). An increase in the frequency of sIPSCs following diazepam suggests enhanced presynaptic GABA release onto DA neurons. Changes in sIPSC amplitude, in contrast, would suggest adaptations in DA neurons, yet diazepam did not alter sIPSC amplitude across all phases of learning (Extended Data Figs. 2D and 2E).

In contrast to lateral shell projections, DA neurons projecting to medial shell and core (Figs. 2G and 2J) showed no significant learning mediated changes in the percentage of sIPSCs with interevent intervals < 10 ms (Figs. 2H and 2K) or total number of spontaneous events (Extended Data Figs. 2F and 2H). Further, bath application of diazepam led to a consistent decrease in sIPSC frequency (Figs. 2I and 2L) without changes in sIPSC amplitude across all stages of learning (Extended Data Figs. 2G and 2I). Overall, these findings highlight a striking circuit specificity of learning-induced KCC2 downregulation, with its impact on VTA GABA neurons selectively innervating DA projections to the NAc lateral shell—a DA pathway that was previously shown to develop phasic cue-evoked responses over the course of learning^8,9^.

### VTA GABA neurons and KCC2 downregulation contribute to reward learning

Given that changes in VTA GABA neurons developed during the acquisition phase of Pavlovian conditioning, we determined whether these neurons mediate cue-reward association. First, we used optogenetics to suppress GABA neuron activity during cue and reward presentations throughout a 13-day Pavlovian training period. GAD-Cre rats were injected with Cre-dependent archaerhodopsin (Arch) and implanted with optic fibers unilaterally in the lateral VTA (Fig. 3A and Extended Data Fig. 3A). When compared to control rats (GFP-expressing with light stimulation and Arch-expressing with no light stimulation, Extended Data Fig. 3B), photoinhibition of VTA GABA neurons in Arch-expressing rats significantly attenuated conditioned responding to the reward-predictive CS (Fig. 3B). No sex differences were observed in conditioned responding (Extended Data Fig. 3C).

**Figure 3.**
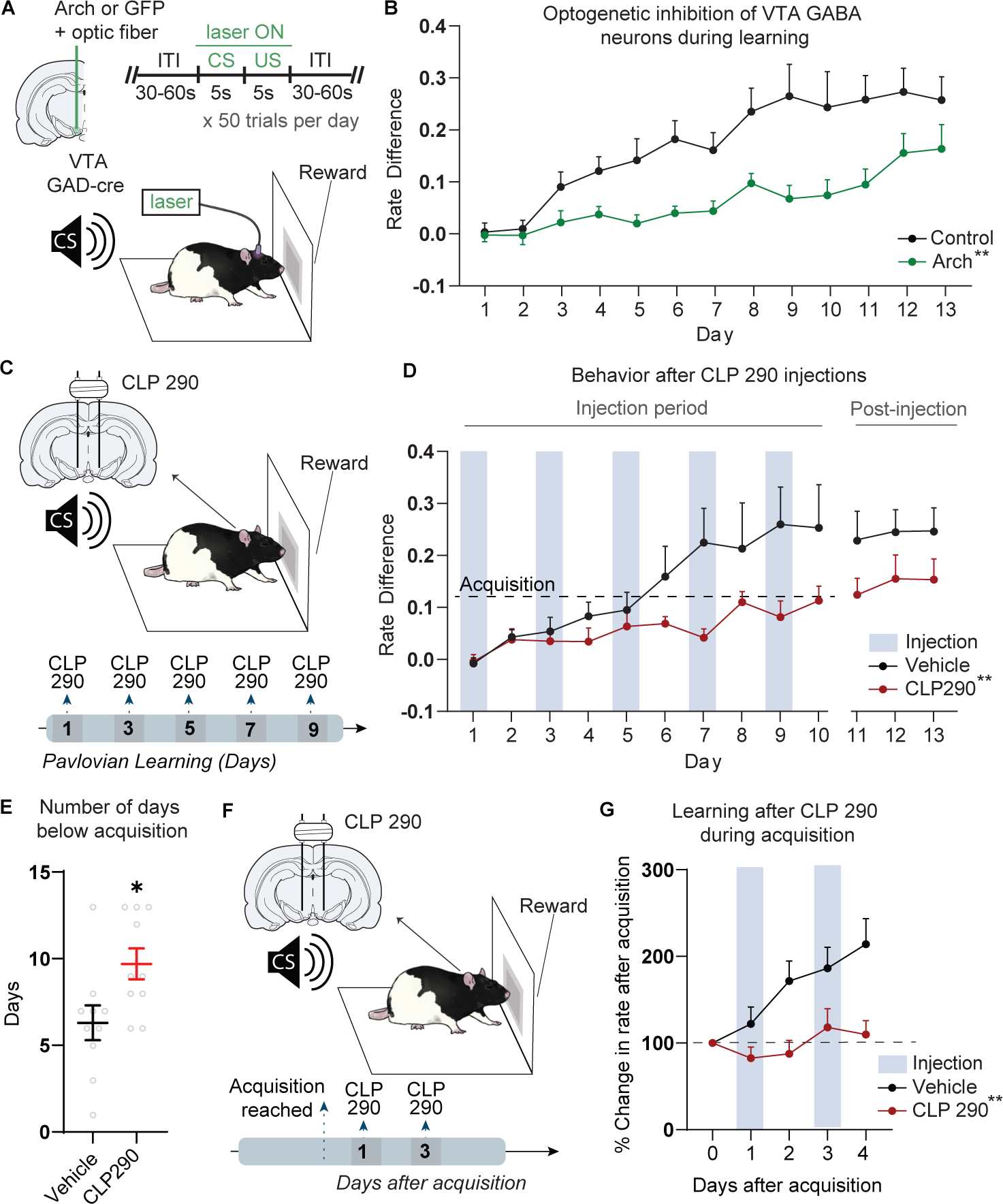
VTA GABA neurons and KCC2 downregulation contribute to reward learning. A) GAD-Cre rats were injected with Cre-dependent Arch in the lateral VTA, followed by the implantation of an optic fiber. During Pavlovian conditioning, green light (520 nm) was delivered throughout the CS and US. B) Green light delivery in Arch-expressing rats attenuated conditioning responding compared to controls. **p < 0.01, RM ANOVA, group x time: F(12, 396) = 2.627, n = 16-19. C) Animals received bilateral intra-VTA infusions of CLP290 or vehicle on alternate days 1 hour before the conditioning session. D) Animals injected with CLP290 on alternate days showed attenuated conditioned responding compared to animals treated with vehicle. The rate difference at the middle of acquisition phase (dashed line) was calculated as half of the rate difference during plateau. **p < 0.01, RM ANOVA, group x time: F(9, 162) = 2.704, n = 10. E) Animals injected with CLP290 spent more days (9.7 ± 2.83%) below the middle of acquisition (defined in D) compared to the animals treated with vehicle (6.3 ± 3.16%). *p < 0.05, t-test, n = 10. F) The effect of bilateral intra-VTA CLP290 administration was assessed starting the day after animal’s response rates reached the middle of acquisition calculated in D. G) In contrast to vehicle, intra-VTA CLP290 administration during acquisition phases suppressed successive learning. **p < 0.01, RM ANOVA, group: F(1, 18) = 13.00, n = 9-11.

Next, we tested whether pharmacological KCC2 activation in the VTA attenuated the formation of cue-reward association. CLP290 (1 µL, 60 µM) or vehicle were bilaterally administered into the lateral VTA (Extended Data Fig. 3D) prior to conditioning sessions on alternate days (Figs. 3C and 3D, shaded blue bars). Compared to rats that received intra-VTA infusion of vehicle (Fig. 3D, black data), intra-VTA infusion of CLP290 significantly decreased conditioned responding to the reward-predictive CS (Fig. 3D, red data). Decreased conditioned responding after CLP290 suggests that preventing KCC2 downregulation in the VTA delays acquisition of cue-reward association. To test this possibility, we calculated the number of conditioning sessions until the animals reached the middle of acquisition phase, defined as half of the rate difference values observed during the plateau phase (dashed line in Fig. 3D). Compared to vehicle-treated controls, CLP290-treated rats had a significantly greater number of training sessions below the middle of acquisition phase (Fig. 3E), suggesting that preventing intra-VTA KCC2 downregulation delayed reward-related learning.

Finally, we examined whether pharmacological KCC2 activation *during* acquisition of Pavlovian conditioning arrested the development of cue-reward association. CLP290 was microinfused bilaterally in the VTA starting the day after animals reached the middle of acquisition phase of learning (Fig. 3F and Extended Data Fig. 3E). In marked contrast to the vehicle-treated rats, which exhibited an increased rate difference, CLP290-treated rats failed to show an enhancement in conditioned responding (Fig. 3G). Taken together, these results indicate that VTA KCC2 downregulation is critical for Pavlovian reward-related learning.

### VTA GABA neurons show enhanced synchronized activity during acquisition periods of learning

We next investigated the mechanisms by which KCC2 downregulation in the VTA contributed to learning. Patch clamp recordings indicated that KCC2 downregulation during acquisition increased the number of concurrent GABA release events with synchrony of less than 10 milliseconds (Figs. 2D and 2E). This phenomenon could arise from increased temporal coincidence of action potential firing between individual GABA neurons in the VTA. To determine whether VTA GABA neurons exhibit millisecond timescale synchrony during Pavlovian conditioning, we performed tetrode and silicon probe electrophysiological recordings in freely behaving rats (Fig. 4A). For cell-type specific recordings, we expressed Cre-dependent Arch and Channelrhodopsin (ChR2) in GAD-Cre and TH-Cre rats, respectively (Extended Data Fig. 4A). Animals were chronically implanted with microdrives carrying recording electrodes and optic fiber cannulas allowing for the classification of lateral VTA cells as putative GABA and DA neurons based on opto-tagging and spike waveform features (Extended Data Figs. 4B and 4C, see Methods).

**Figure 4.**
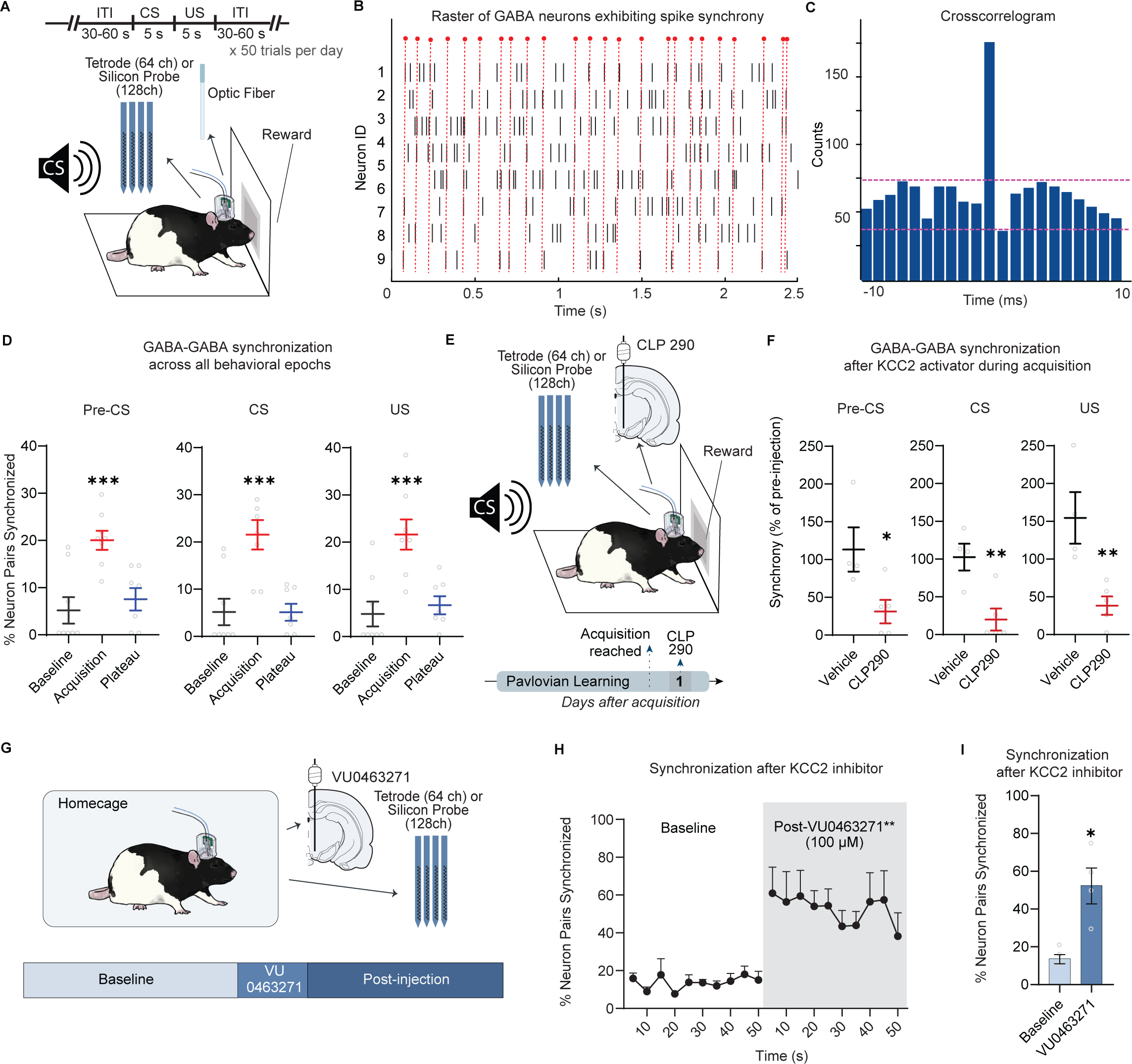
VTA GABA neurons show enhanced coordinated activity during acquisition periods of learning. A) A microdrive carrying an optic fiber and 16 tetrodes (or 128-channel silicon probe) was unilaterally implanted in TH-Cre and GAD-Cre for cell-type specific recordings in the lateral VTA during Pavlovian conditioning (see methods). B) Representative raster of putative VTA GABA neurons across time. The vertical red dashed lines indicate instances of synchronized firing between two or more neurons. C) Representative cross-correlogram between a pair of putative GABA neurons exhibiting millisecond timescale synchrony. Magenta lines indicate global significance bands. D) The mean percent of putative GABA pairs synchronized across each phase of learning during intertrial interval (pre-CS), cue (CS) and reward (US) presentation. Synchronization was enhanced during acquisition periods (red) of learning compared to baseline and plateau. ***p < 0.001, one-way ANOVA, F = 11.18 (pre-CS), F = 12.52 (CS), and F = 12.06 (US), n = 7-8 rats. E) Microdrives coupling tetrodes or silicon probes with microinfusion cannulas were implanted for intra-VTA electrophysiological recordings and local drug administration. The effect of unilateral intra-VTA CLP290 administration was assessed the day after the animal’s response rates reached the middle of acquisition (calculated in S4D). F) Intra-VTA CLP290 administration during acquisition significantly decreased the normalized number of synchronized GABA pairs compared to the vehicle-treated groups across pre-CS, CS, and US. The numbers of synchronized putative GABA pairs were normalized to the day preceding injections. *p < 0.05, **p < 0.01, t-test, n = 4-5 rats. G) Tetrode or silicon probe recordings were combined with unilateral intra-VTA microinfusions of VU0463271 in naïve animals within their homecages. VTA GABA synchronization was analyzed over a 50-s time period before and after VU0463271 microinfusion. H) VTA GABA neuron synchronization during 50-second periods before and after VU0463271 administration. VU0463271 significantly increased the number of synchronized GABA pairs compared to the pre-infusion baseline. **p < 0.01, RM ANOVA, group: F(1, 6) = 15.67, n=4.I) The percentage of synchronized GABA neuron pairs averaged across 50-seconds significantly increased after VU0463271 administration compared to the pre-infusion baseline. *p < 0.05, t-test, n = 4.

In agreement with previous work, the proportion of lateral VTA DA and GABA neurons responding to CS increased with the progression of associative learning^14,15^ (Extended Data Figs. 4D-4G). At all stages of learning, we observed synchronous spiking between individual VTA GABA neurons (Fig. 4B). To quantify changes in synchrony across learning phases, we calculated cross-correlograms (CCGs) between pairs of simultaneously recorded putative lateral VTA GABA neurons^34,35^ (Fig. 4C, see Methods). Our results reveal that during acquisition, a significantly higher percentage of putative GABA neuron pairs exhibited millisecond timescale synchrony compared to baseline and plateau phases of learning (Fig. 4D). Importantly, significant increases in synchrony were observed beyond the presentation of the CS and US, encompassing the intertrial intervals preceding them (Fig. 4D, Pre-CS).

We then postulated that pharmacological manipulation of KCC2 in the VTA would alter synchronized activity of VTA GABA neurons *in vivo*. Rats were chronically implanted with a microdrive system designed for unilateral drug infusions, opto-tagging, and simultaneous electrophysiological recordings from the lateral VTA (Fig. 4E). First, we microinfused KCC2 activator CLP290 (1 µL, 60 µM) or vehicle in the VTA of learning animals once they reached the middle of the acquisition period (defined based on animal’s performance in Extended Data Fig. 4D). Across all parts of a trial, lateral VTA GABA neurons exhibited a significant decrease in coordinated firing activity after intra-VTA CLP290 injections compared to vehicle treatment (Fig. 4F). Importantly, intra-VTA CLP290 injections did not impact locomotion (Extended Data Figs. 5A-5C). Next, we microinfused KCC2 antagonist VU0463271 (1 µL, 100 µM) in the lateral VTA of naïve, freely moving animals in their home cage (Fig. 4G). When compared to the pre-injection period, VU0463271 significantly increased the mean percentage of synchronized VTA GABA neuron pairs (Figs. 4H and 4I). All probe implantation and microinfusion sites in the VTA are shown in Extended Data Fig. 4H. Cumulatively, these findings indicate that during acquisition phases of reward learning, individual VTA GABA neurons synchronize their activity within a millisecond time frame, a phenomenon bidirectionally dependent on KCC2 function.

### Changes in KCC2 function enhance DA signaling in the NAc and VTA

Since reward learning involves DA signaling in the NAc lateral shell^9^, we hypothesized that KCC2 downregulation in the VTA might also alter DA release in this region. To test this hypothesis, we injected a genetically encoded DA sensor GRAB_DA2m_^36^ and implanted an optical fiber in the NAc lateral shell (Extended Data Fig. 6A and Fig. 5A). Rats were subjected to the Pavlovian conditioning task and DA transients were measured during baseline and acquisition of learning (Fig. 5A). Fiber photometry recordings in the NAc revealed increased DA responses to the reward-predictive cue, while DA responses to reward delivery remained unchanged (Fig. 5B and Extended Data Fig. 6B). Unilateral CLP290 microinfusions the day after animals reached the middle of acquisition (Fig. 5C) attenuated DA responses to CS in the lateral shell with no changes in DA responses to the US (Figs. 5D and 5E).

**Figure 5.**
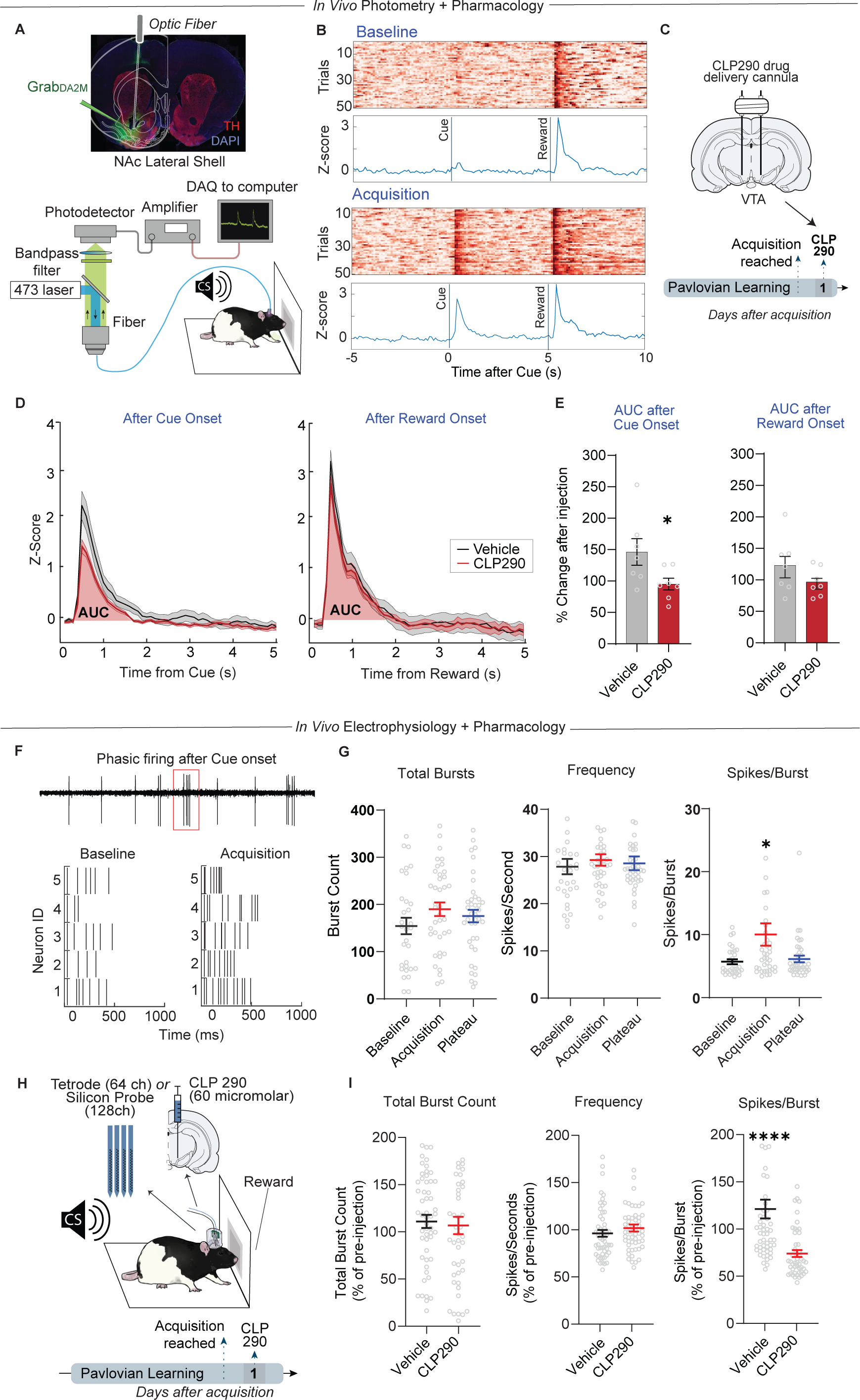
Changes in KCC2 function enhance DA signaling in the NAc and VTA. A) Top: GRAB_DA2m_ (green) was expressed unilaterally in the NAc lateral shell, followed by an optic fiber implantation. TH-expressing DA terminals are shown in red. Bottom: Schematics showing fiber photometry recording of GRAB_DA_ sensor recordings while the animal undergoes Pavlovian training. B) Representative heatmaps for GRAB_DA2m_ in the NAc lateral shell showing individual cue and reward responses (50 trials) during baseline and acquisition. Below are the average z-scored traces across trials. C) Fiber photometry recordings were combined with CLP290 or vehicle intra-VTA microinfusions the day after the middle of acquisition was reached. D) Comparison of Z-score averages for GRAB_DA_ responses to cue and reward in animals treated with CLP290 (red) or vehicle (blue). Area under the curve (AUC, shaded) of the signals were calculated for averaged trials. E) The mean changes in DA release (quantified as AUC) during cue (left) and reward (right) presentations after CLP290 and vehicle administration. AUCs on the day of infusions were normalized and expressed as percent change relative to the previous day. CLP290 administration significantly reduced DA responses to cues but not to rewards (p = 0.19). *p < 0.05, t-test, n = 7. F) Representative spike raster from DA neurons engaging in phasic firing after cue presentation (top). Raster plots of isolated burst trains during baseline and acquisition (bottom). G) Burst firing parameters in DA neurons were assessed during CS across baseline, acquisition, and plateau phases of learning. Across all stages, DA neurons exhibited no significant differences in total burst count (left, p = 0.68) and intra-burst frequency (middle, p = 0.27) during CS. In contrast, the number of spikes per CS-evoked burst was significantly increased during acquisition (10.14 ± 1.62%) compared to baseline (5.8 ± 1.04%) and plateau (6.24 ± 0.99%). *p < 0.05, one-way ANOVA, F = 4.63, n = 26-40 units, 5-8 rats. H) Lateral VTA neuron activity was assessed with tetrode or silicon probe recordings and CLP290 was unilaterally infused in the VTA on the day after animal’s response rates reached the middle of acquisition (calculated in Figure 4D). I) Average CS-evoked burst firing parameters after CLP290 and vehicle administration were normalized and expressed as percent change relative to the pre-administration day. CLP290 microinfusion produced no changes in total burst (left, p = 0.72) and frequency (middle, p = 0.28), but significantly reduced the number of spikes per burst compared to vehicle-infused group (right). ****p < 0.0001, t-test, n = 49-55 units, 4-5 rats.

Given that reward-predictive cues stimulate DA release by increasing phasic firing of lateral VTA DA neurons^8,37^, we determined whether KCC2 downregulation altered cue-evoked DA neuron bursting activity during acquisition of learning. First, we assessed the burst firing parameters of putative VTA DA neurons during CS across all learning stages in animals with tetrode and silicon probes (Fig. 5F). No significant differences in total burst count during CS and mean spike frequency within a burst were detected between baseline, acquisition, and plateau phases of learning (Fig. 5G, left and middle). However, the mean number of spikes within a burst was significantly higher during acquisition (Fig. 5G, right). Next, we reassessed the burst firing parameters during CS in animals administered with intra-VTA CLP290 or vehicle the next day after reaching the middle of acquisition (Fig. 5H). We did not observe significant differences in the total number of bursts and intra-burst spike frequency during CS in VTA DA neurons between CLP290 and vehicle-treated groups (Fig. 5I, left and middle). However, after CLP290 injections, DA neurons exhibited a significant decrease in the mean number of spikes within bursts compared to vehicle treatment (Fig. 5I, right).

### Optogenetic synchronization of VTA GABA neurons enhances stimulus-induced burst firing *ex vivo* and *in vivo*

Based on our findings in Figs. 4 and 5, we hypothesized that VTA GABA network synchronization potentiates phasic DA firing. To test this, we aimed to mimic learning-mediated neuronal synchronization by using optogenetic stimulation of VTA GABA neurons. First, we determined the frequency at which spikes in one VTA GABA neuron exhibited millisecond synchrony with spikes in another VTA GABA neuron during the acquisition period. To this end, we generated joint peristimulus time histograms that enabled the quantification of time intervals between two consecutive instances of synchrony in spike trains from pairs of VTA GABA neurons (Extended Data Fig. 7A). This analysis revealed that during acquisition, pairs of VTA GABA neurons displayed millisecond time frame synchrony at a mean frequency of around 10-Hz (Extended Data Figs. 7B and 7C).

Next, we studied the effects of optogenetically-induced GABA network synchronization on DA bursting. GAD-Cre animals were injected with Cre-dependent ChR2 in the VTA to stimulate GABA neurons at 10-Hz frequencies. In the first set of experiments, we prepared midbrain horizontal slices and performed patch-clamp cell-attached recordings of DA neurons (Fig. 6A). Putative DA neurons in the lateral VTA were identified using established electrophysiological criteria and suppression of spontaneous action potential firing during ChR2 stimulation (Extended Data Figs. 8A-8D, see Methods). Phasic bursts of action potentials were triggered by direct iontophoretic application of glutamate near recorded neurons. Consistent with previous reports, iontophoretic application of glutamate elicited high-frequency spike trains in lateral VTA DA neurons (Fig. 6B). Strikingly, the same DA neurons showed an increased number of spikes per burst when glutamate application was paired with 10-Hz light-induced synchronization of VTA GABA neurons (Figs. 6C and 6D). Optogenetic VTA GABA neuronal synchronization did not alter the mean spike frequency within glutamate-induced bursts in DA neurons (Extended Data Fig. 8E).

**Figure 6.**
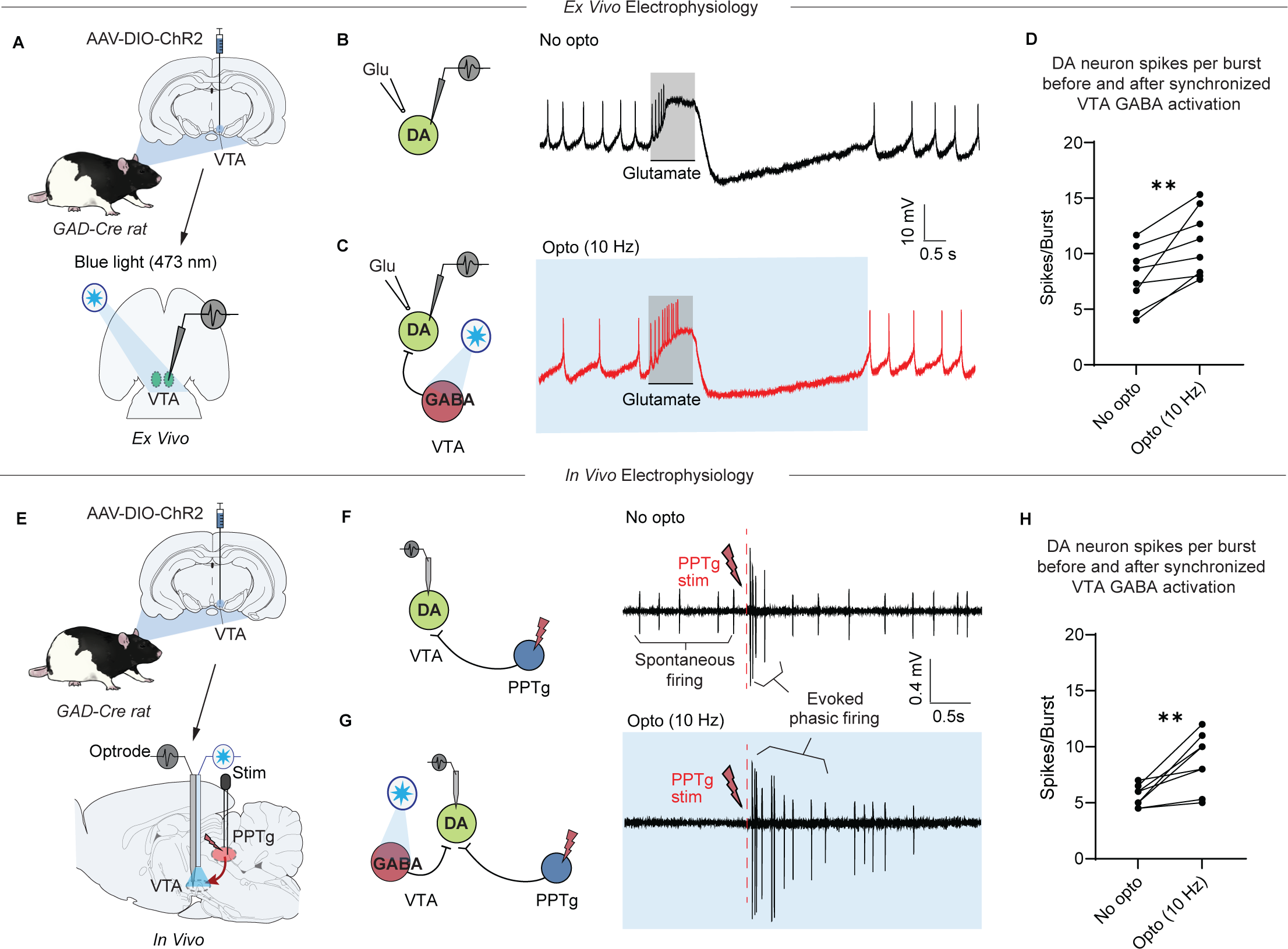
Optogenetic synchronization of VTA GABA neurons enhances stimulus-induced burst firing *ex vivo* and *in vivo*. A) Cre-dependent ChR2 was expressed in the VTA of GAD-Cre animals. Two-to-three weeks after surgery, cell-attached patch clamp recordings of lateral VTA DA neurons were combined with optogenetic synchronization of VTA GABA neurons in midbrain horizontal slices. B) Representative trace showing that in spontaneously active lateral VTA DA neurons, iontophoretic application of glutamate (Glu) triggers high-frequency burst firing. For display, the traces were filtered. C) In the same DA neuron (from B), light-induced synchronized activation of VTA GABA neurons at 10-Hz frequency facilitated glutamate-induced phasic firing (increased spikes within bursts). D) Light-induced VTA GABA neuron synchronization significantly increased the number of spikes within glutamate-evoked bursts in VTA DA neurons. **p < 0.01, paired t-test, n = 8 cells, 3 rats. E) Cre-dependent ChR2 was injected in the VTA of GAD-Cre rats. Two-to-three weeks after surgery, single-unit recordings in anesthetized rats were paired with optogenetic stimulation of VTA GABA neurons (optrode). Phasic DA firing was evoked by electric stimulation of excitatory afferents from the pedunculopontine tegmentum (PPTg, Stim). F) Representative trace showing that PPTg stimulation elicited a phasic burst in a spontaneously active lateral VTA DA neuron. For display, traces were filtered and stimulus artifacts were removed. G) In the same cell, PPTg stimulation was paired with optogenetic 10-Hz synchronized activation of VTA GABA neurons. Light-induced VTA GABA synchronization attenuated spontaneous tonic firing, yet also potentiated PPTg-driven phasic bursting in DA neurons (increased spikes within bursts).H) Light-induced 10-Hz VTA GABA neuron synchronization significantly increased the number of spikes within PPTg-stimulated bursts in lateral VTA DA neurons *in vivo*. **p < 0.01, paired t-test, n = 8 cells, 5 rats.

Qualitatively similar effects of GABAergic synchronization on phasic DA firing were observed during *in vivo* single-unit recordings of VTA DA neurons in anesthetized rats. Two-to-three weeks after Cre-dependent ChR2 injections, a glass electrode coupled to an optic fiber was lowered to the lateral VTA of GAD-Cre rats (Fig. 6E). Putative lateral VTA DA neurons were identified based on their electrophysiological properties and light-induced suppression of spontaneous action potential firing (Extended Data Figs. 8F-8G, see Methods). Stimulus-evoked burst activity in DA neurons was assessed via electric stimulation of the pedunculopontine tegmental nucleus (PPTg, Fig. 6E), which relays cue-related sensory information to these cells^38–40^. In 8 of 15 recorded DA neurons, PPTg stimulation elicited a burst of action potentials (Fig. 6F). In those cells in which bursts were evoked, PPTg stimulation was repeated with concurrent optogenetic synchronization of VTA GABA neurons at 10-Hz. Notably, optogenetic VTA GABA synchronization potentiated PPTg-driven phasic bursting by increasing the number of spikes within a burst (Figs. 6G and 6H). In contrast, synchronizing GABA neuron activity did not alter the intraburst firing rates in lateral VTA DA neurons (Extended Data Fig. 8H).

## Discussion

Associative reward learning is pivotal for an organism’s survival, but the neuronal adaptations underlying this fundamental process have not been well delineated. We found that circuit-specific changes in Cl^−^ homeostasis within GABA neurons of the VTA contribute to the formation of Pavlovian cue-reward association. Specifically, during the acquisition phase of learning, neuron-specific Cl^−^ transporter KCC2 undergoes transient downregulation, manifesting as depolarized E_GABA_ in VTA GABA neurons. These alterations were associated with increased firing synchrony within VTA GABA neuronal networks and enhanced phasic bursting in VTA DA neurons. Most importantly, enhancing KCC2 function or silencing GABA networks in the VTA during acquisition of learning attenuated the formation of cue-reward associations.

Despite considerable focus on VTA DA neurons in reward learning^41,42^, learning-induced alterations in VTA GABA neurons remain poorly understood. To address this gap, we used a photoconvertible Ca^2+^ indicator to focus exclusively on VTA GABA cell populations that were active during cue and reward presentations. In these populations, Cl^−^ homeostasis was altered only during the acquisition phase of learning, when animals exhibited the largest change in conditioned responding. Furthermore, we found post-translational downregulation of functional KCC2 (Fig. 1J) and impaired GABA_A_ receptor-mediated inhibition (Fig. 2F) during acquisition, but not maintenance of cue-reward association. Interestingly, similar transient dynamics were observed in regard to learning-induced changes in glutamatergic receptors on DA neurons^42^, suggesting that transient changes in VTA neurons act to establish new associations but are not necessary for the long-term storage of cue-reward information. The preservation of reward-related memories may rely on other mechanisms outside the VTA, and restoring Cl^−^ homeostasis after acquisition of associations would permit future reward learning^40^.

Prior studies showed that KCC2 downregulation causes paradoxical excitation of VTA GABA neurons in response to GABA_A_ receptor stimulation^17,20^. Perhaps unsurprisingly, when we enhanced GABA_A_ receptor function in VTA GABA neurons with diazepam, reduced KCC2 function led to increased GABA release on downstream targets (Fig. 2F). The unexpected finding, however, was that during acquisition phases of learning, KCC2 downregulation increased coincident spiking amongst VTA GABA neurons (Figs. 2E and 4). While previous studies have noted enhanced coordination of midbrain activity after learning^43,44^, our findings show that the acquisition of conditioned learning is disrupted when network synchronization is suppressed by KCC2 activation. The relationship between reduced KCC2 activity and neuronal synchronization in the adult brain has been previously associated with pathological conditions^45,46^. However, growing evidence implicates Cl^−^ transport changes in neuronal synchronization in the context of seasonal and sleep-related variations observed in the hypothalamus and cortex^47,48^. Integrating our findings with the existing literature, we propose that changes in KCC2 may be a common mechanism underlying neuronal synchronization under physiological conditions in the adult brain.

At first glance, the notion that KCC2 downregulation and resulting GABAergic synchronization enhance cue-induced phasic DA signaling might appear contradictory. Previous work has associated KCC2 downregulation with reduced DA neuron firing in the VTA and blunted DA release in the NAc^17,20,21^; however, these studies primarily examined DA firing in acute brain slices or anesthetized animals. Additionally, measurements of DA release in the NAc were conducted using microdialysis over minutes, which may not capture millisecond-scale changes in phasic DA release. Despite reports of attenuated DA activity, previous studies have also identified correlations between KCC2 downregulation in VTA GABA neurons and increased consumption of addictive drugs^17,20^. Hence, we propose that enhanced phasic DA signaling due to KCC2-induced changes in VTA GABA neuron firing patterns may constitute one mechanism that promotes addictive behaviors. In support of this hypothesis, mounting evidence demonstrates that VTA GABA neurons exhibit excitatory responses in response to reward-related stimuli^34,49^ (Extended Data Fig. 4). There is also increasing recognition of the crucial role of concerted activity in both VTA DA and GABA neurons in enhancing phasic DA neuron firing^34,50^. Most notably, mathematical simulations revealed that synchronized inhibitory input to DA neurons can evoke additional spikes during bursting^34^. Moreover, electrophysiological experiments suggest that brief inhibitory pulses interleaved with depolarizing currents can extend burst firing in midbrain DA neurons^51^. These studies suggest two major mechanisms by which synchronized inhibition can augment DA burst responses: First, removal of inhibition between GABAergic pulses increases the probability of spike generation in DA neurons^34^. Second, GABA-mediated hyperpolarization attenuates depolarization-induced inactivation of the sodium current in DA neurons, a putative mechanism for burst firing termination^51^. Our slice and *in vivo* experiments strongly support these mechanisms by showing that optogenetic synchronization of VTA GABA neurons enhances phasic bursting of DA neurons via increasing the number of spikes within a burst (Fig. 6).

Our findings demonstrate a notable circuit-specificity within the mesolimbic system, wherein learning-dependent KCC2 downregulation exclusively impacts inhibitory transmission to DA neurons that project to the NAc lateral shell. Extensive evidence indicates that separate DA projections to distinct NAc subregions mediate different aspects of reward processing. Our results add to the body of work showing that NAc lateral shell projecting DA neurons exhibit sharp activity increases to reward-relevant stimuli, and further highlight the importance of lateral mesolimbic DA pathways in reward-related learning^8,9^. Although much less is known about the interaction between heterogeneous DA populations and their inhibitory neighbors, some results have shown that lateral NAc projections receive a considerably stronger local inhibitory input compared to medial NAc projection^52^. Thus, the lack of KCC2 downregulation on NAc medial shell and core-projecting DA neurons may arise from simply a difference in the number of synapses from VTA GABA neurons. Nevertheless, we do not rule out the possibility that medial subsets of DA neurons are contacted by distinct subclasses of VTA GABA neurons.

A guiding principle for the development of neuropsychiatric disorders is that normal learning and memory processes are ‘usurped’ into a pathological state^53^. Several studies have highlighted that KCC2 downregulation represents a common yet often overlooked form of VTA GABA-specific dysregulation in maladaptive states^17,18,20^. Our results suggest that this GABA-specific switch in Cl^−^ gradient is a normal form of neuroadaptation during learning that is ‘hijacked’ in addiction and other neuropsychiatric disorders. In summary, given that impaired reward learning is central to many neuropsychiatric disorders and aberrant function of KCC2 is observed after exposure to drugs of abuse and stress, our findings provide a new context for understanding neuropsychiatric disorders and offer a rich avenue for future investigations.

## Acknowledgements

We thank Dr. Patrick Forcelli, Dr. Stefano Vicini, Dr. Daniil Berezhnoi, and Anna Pearson for helpful discussions and feedback. This work was supported by grants from the National Institutes of Health [MH125996 (AO), DA048134(AO), NS139517 (JW), DA061493 (JW)], Brain & Behavior Research Foundation [NARSAD 28113 (AO)], and the Whitehall Foundation [2020-12-35 (AO)].

## Contributions

J.W. and A.U. designed and performed ex vivo and in vivo electrophysiological experiments assisted by I.C. and A.K.S. D.R. and C.C.S performed behavioral experiments. J.W., A.U. and H.C.S. performed western blot and immunohistochemistry experiments. A.O. originated, planned, and oversaw the experiments with J.W.’s and A.U.’s assistance. Led by A.O., all the authors contributed to writing the manuscript.

