## Supplemental Data for "Dynamic Changes in Chloride Homeostasis Coordinate Midbrain Inhibitory Network Activity during Reward Learning"

Supplemental Figures (S1-S8)

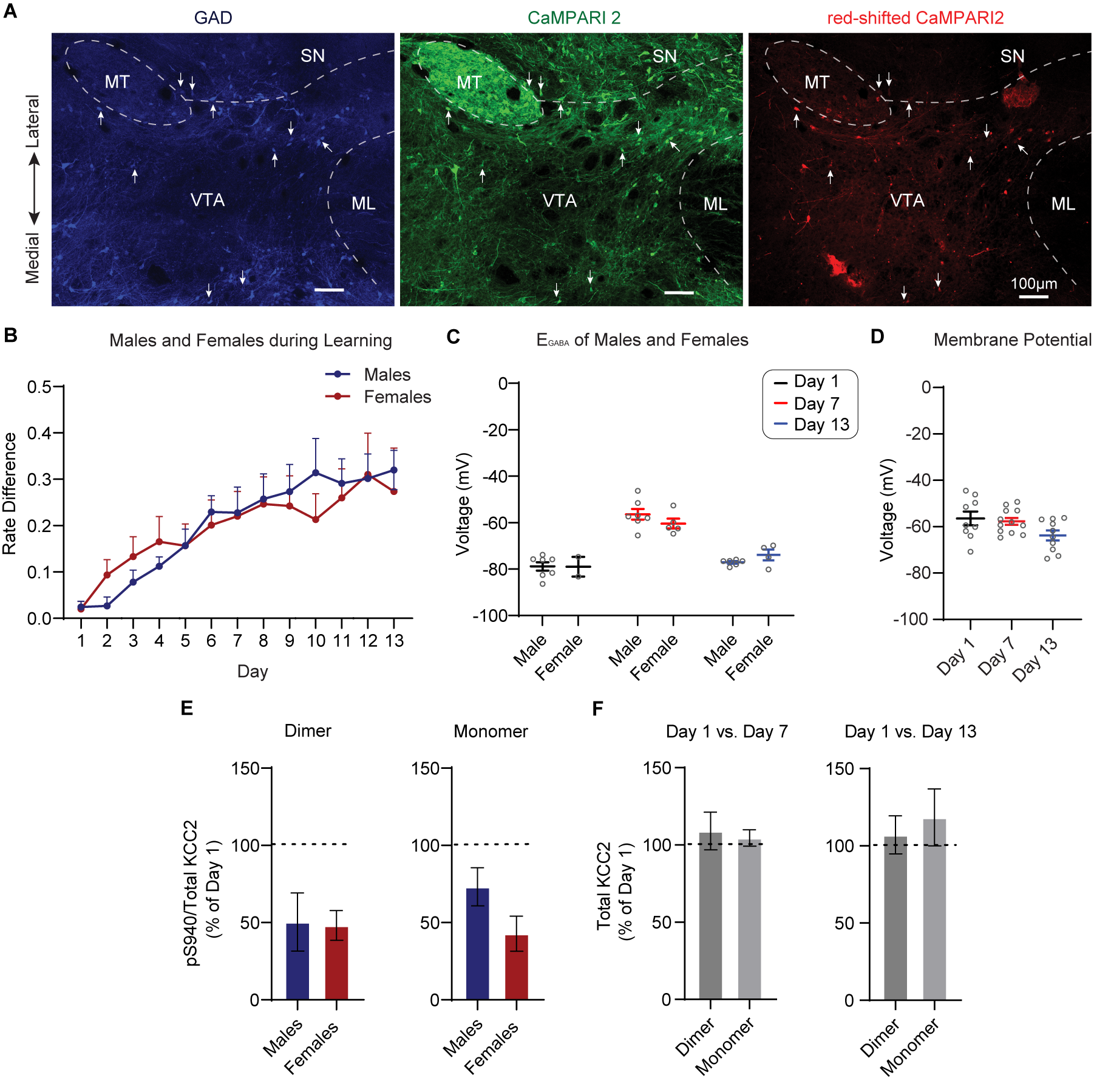
**Figure S1. Learning-dependent downregulation of functional KCC2 in male and female rats. Related to Figure 1.**

A) Immunohistochemical labeling of CaMPARI2-expressing VTA GABA neurons and red-shifted CaMPARI2-expressing neurons, as detailed in the methods. Blue fluorescence highlights VTA GABA neurons with BFP, while green fluorescence indicates CaMPARI2 expression. Red fluorescence identifies photo-converted CaMPARI2 expression. Arrows indicate neurons exhibiting fluorescence for all three channels.

B) Behavioral trajectory of males and females did not differ significantly across time (p = 0.95, sex x time: F (1, 37) = 0.02, n = 12-22 rats, two-way ANOVA).

C) E_GABA_ values did not exhibit significant differences between sexes (p = 0.23, sex x time: F (2,25) = 1.56, n = 2-9 rats, two-way ANOVA)

D) Resting membrane potentials of the paired learning group did not show significant differences across time (p = 0.06, F = 3.2, n = 9-12 cells, 5-7 rats, one-way ANOVA)

E) Expression of ps940 KCC2 did not significantly differ between sexes (p =0.92, Dimer; p = 0.15, Monomer, t-test)

F) Expression of total KCC2 protein between Day 1 vs. Day 7 (p = 0.59, Dimer; p = 0.63, Monomer, n = 6, paired t-test) and Day 1 vs. Day 13 (p = 0.72, Dimer; p = 0.7, Monomer, n = 8, paired t-test).

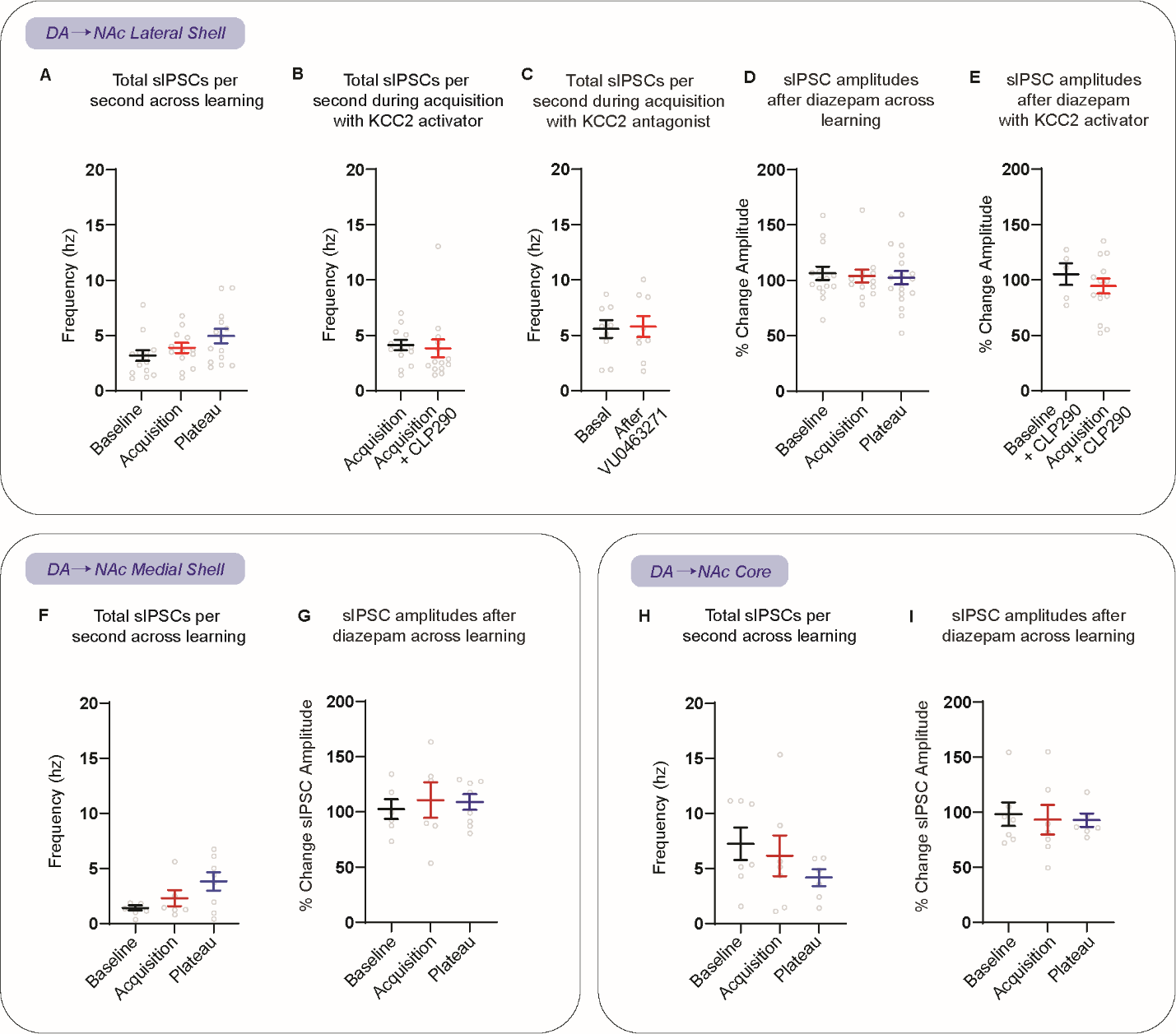

**Figure S2. SIPSC frequencies and amplitudes across mesolimbic DA projections. Related to Figure 2.**

A) The frequency of spontaneous events in NAc lateral shell-projecting DA neurons showed no changes across learning (p = 0.08, F = 2.77, n = 13-14 cells, 3-4 rats, one-way ANOVA).

B) The frequency of spontaneous events in NAc lateral shell-projecting DA neurons showed no changes after CLP290 incubation during acquisition (p = 0.3, n = 13-14 cells, 3-4 rats, Mann-Whitney test).

C) The frequency of spontaneous events in naive animals showed no changes before and after bath application of VU0463271 (p = 0.5, n = 9 cells, 3 rats, paired t-test).

D) The amplitudes of sIPSCs showed no changes after bath application of diazepam across distinct phases of learning in NAc lateral shell-projecting DA neurons (F = 0.69, n = 13-14 cells, 3-4 rats, one-way ANOVA).

E) The amplitudes of sIPSCs showed no changes following bath application of diazepam in CLP290-incubated slices from baseline and acquisition phases of learning (p = 0.41, n = 5-14 cells, 2-4 rats, t-test).

F) The frequency of spontaneous events in NAc medial shell-projecting DA neurons showed no changes across learning (p = 0.09, F = 2.79, n = 6-8 cells, 4 rats, one-way ANOVA).

G) The amplitude of sIPSCs showed no significant changes after bath application of diazepam in NAc medial shell-projecting DA neurons across learning (p = 0.96, F = 0.04, n = 6-8 cells, 4 rats, one-way ANOVA).

H) The frequency of spontaneous events in NAc core-projecting DA neurons showed no changes across learning (p = 0.37, F = 1.47, n = 6-7 cells, 3-5 rats, one-way ANOVA).

I) The amplitudes of sIPSCs showed no significant changes after bath application of diazepam in NAc core-projecting DA neurons across learning (p = 0.85, F = 0.17, n = 6-7 cells, 3-5 rats, one-way ANOVA).

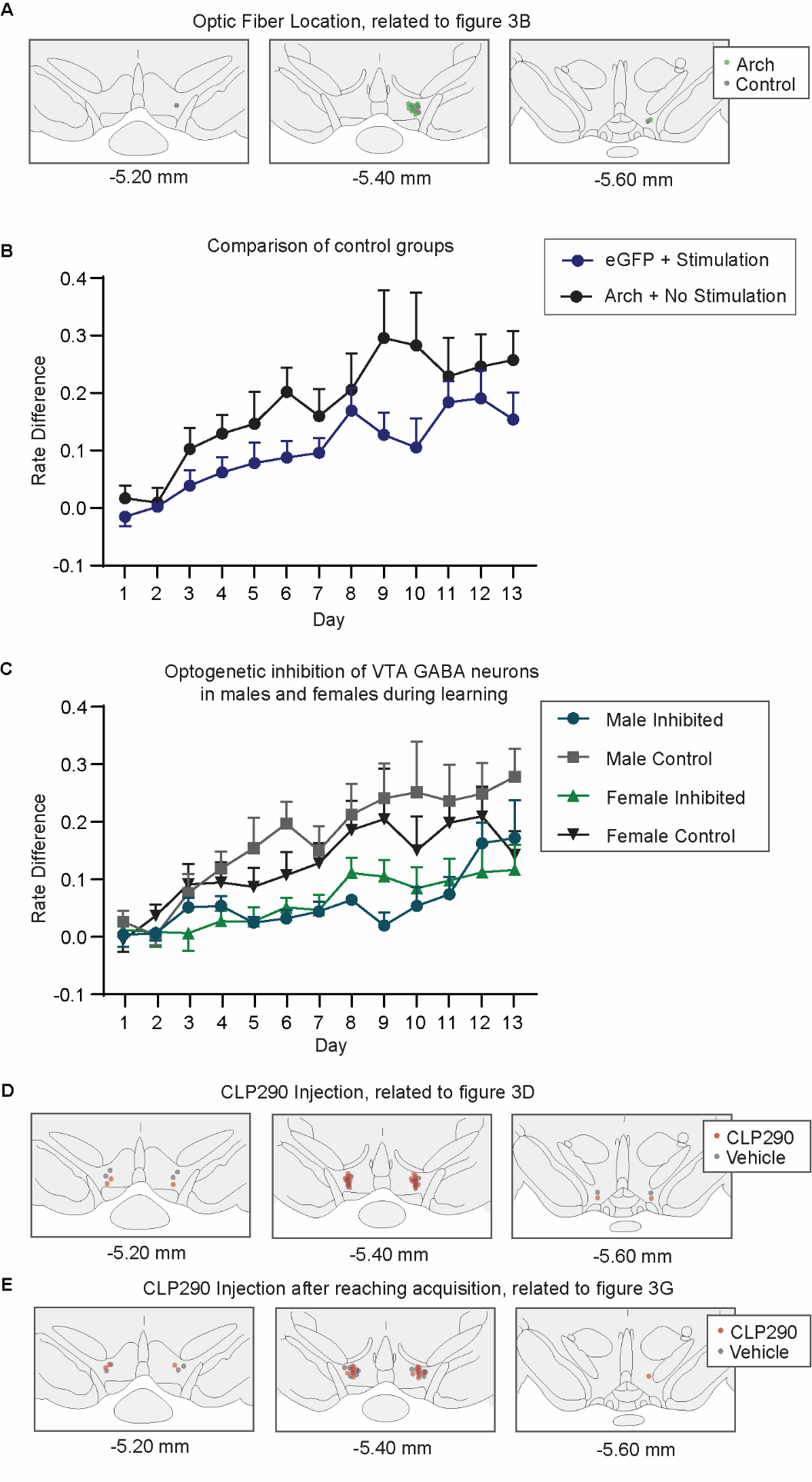

**Figure S3. Optogenetic inhibition of GABA neurons and microinfusions of CLP290 in the VTA during reward learning. Related to Figure 3.**

A) Anatomical placements of the optic fiber probes and Arch injections were determined as described in the methods. The distribution of the optic fiber was similar between groups.

B) No significant differences were observed with the control conditions where eGFP was expressed with light stimulation, and Arch was expressed with no light stimulation (p = 0.1773, time x control groups: F (12, 168) = 1.385, n = 8 rats, two-way RM ANOVA).

C) No significant sex differences were observed in control and opto-inhibition groups (p = 0.47, sex: F (1, 31) = 0.5297, n = 7-10 rats, three-way ANOVA).

C) Anatomical placements of the drug delivery cannula were determined as described in the methods. The distribution of the injection sites was similar between cohorts that received vehicle and CLP290.

D) Anatomical placements of the drug delivery cannula for experiments that entailed CLP290 delivery after acquisition were determined as described in the methods. The distribution of the injection sites was similar between cohorts that received vehicle and CLP290.

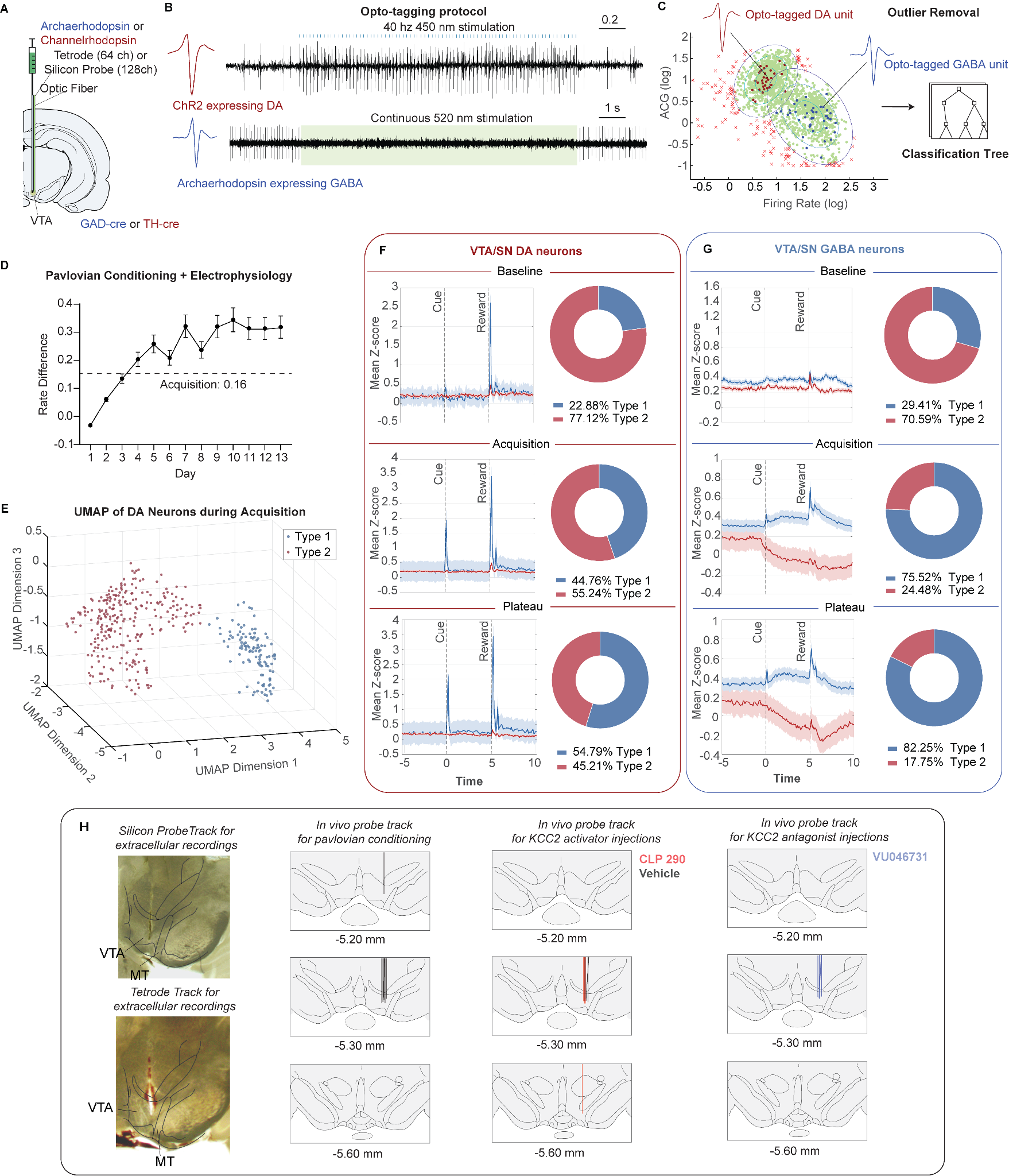

**Figure S4. Cell-type classification in learning rats, and anatomical placement of probes. Related to Figure 4.**

A) Arch was injected in the VTA of GAD-Cre animals, while ChR2 was injected in TH-Cre animals for opto-tagging. Tetrodes or silicon probes were implanted in tandem with an optic fiber (see methods).

B) Top: sample trace for putative DA neuron in a TH-Cre rat showing increased firing rate in response to 40 Hz blue light stimulation. Bottom: example trace for putative GABA neuron in a GAD-Cre animal showing reduced firing rate in response to continuous green light stimulation.

C) Cell type classification. Left panels show two 2-D gaussian mixture models (GMM) fit to the tagged units (DA unit: red, GABA unit: blue). Ellipses show iso-probability regions for each Gaussian. Units within 3 standard deviations from the centroid of each gaussian were utilized for a classification tree (right). See methods.

D) Learning trajectory of animals implanted with probes for extracellular recordings. The rate difference for the middle of acquisition (dashed line) was calculated as half of the rate difference observed during plateau.

E) To characterize the response of DA and GABA populations to cue and reward at each learning stage, the firing rate of putative DA and GABA units was Z-scored. Next, we transformed diverse Z-scored changes in firing rate into a low-dimensional space using Uniform Manifold Approximation and Projection (UMAP). Finally, to classify neuronal responses, we used hierarchical clustering with the Elbow Method to determine the optimal number of clusters within each data set (see methods).

F) Z-scored average firing rate responses to cue and reward, along with the distribution of ‘type 1’ and ‘type 2’ DA neurons during baseline, acquisition, and plateau phases. See methods.

G) Z-scored average responses to cue and reward, along with the distribution of clustered ‘type 1’ and ‘type 2’ GABA neurons during baseline, acquisition, and plateau phases.

H) Left two panels: anatomical placements of probes for extracellular recordings during naturalistic Pavlovian conditioning. Right panels: anatomical placements of the drug delivery cannulae for experiments that entailed CLP290 and VU0463271 microinfusions.

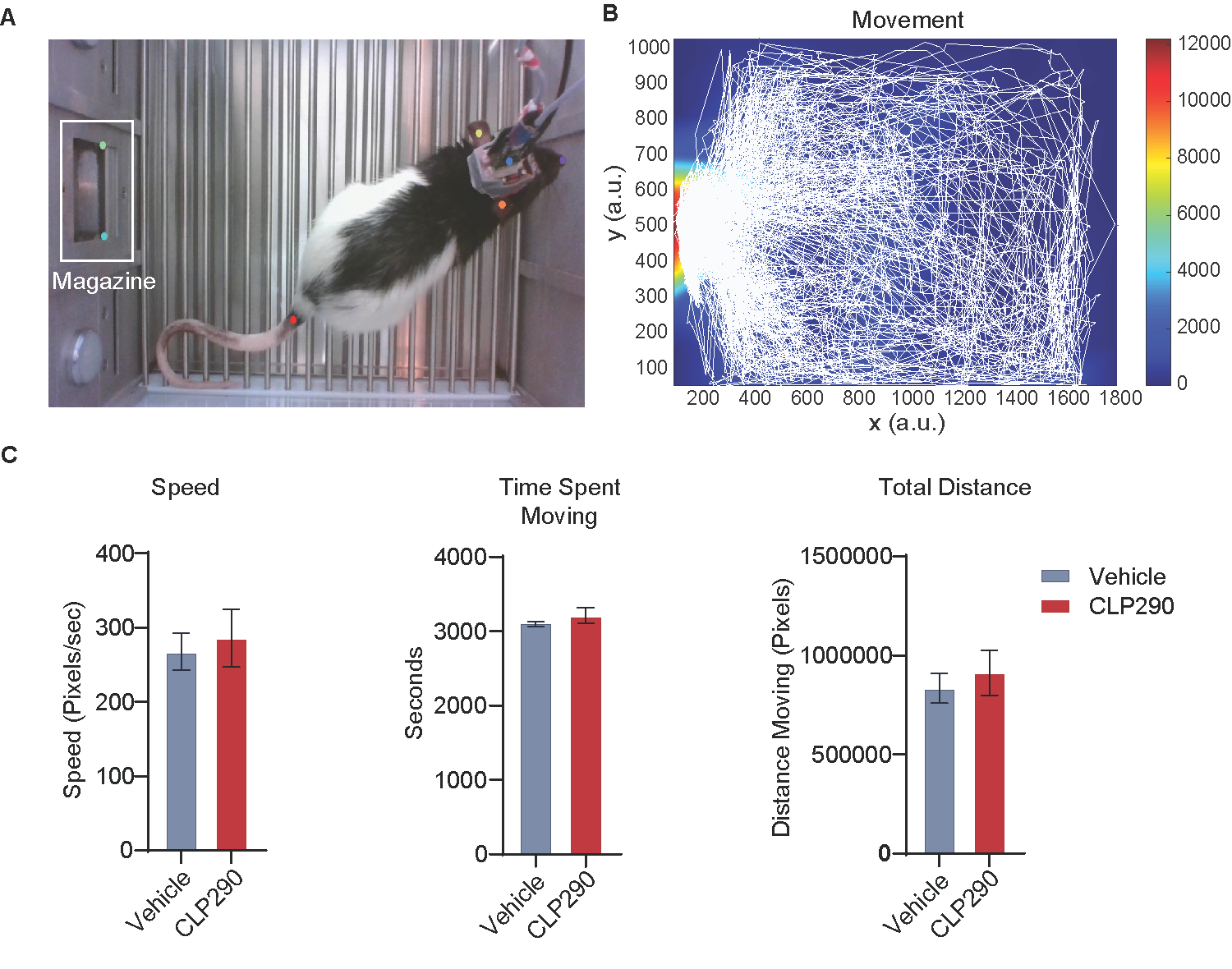

**Figure S5. Intra-VTA CLP290 injections do not impact animal’s locomotor activity. Related to Figure 4.**

A) Representative frame from a video with body parts and magazine (white) labeled from Deep Lab Cut (ears: yellow and orange, headstage: blue, and tail: red). See methods.

B) Representative image of movement tracking during the conditioning paradigm.

C) CLP290 injections did not significantly impact speed (p = 0.99) time spent moving (p = 0.39), and total distance (p = 0.59, n = 4 rats, t-test) compared to vehicle injections.

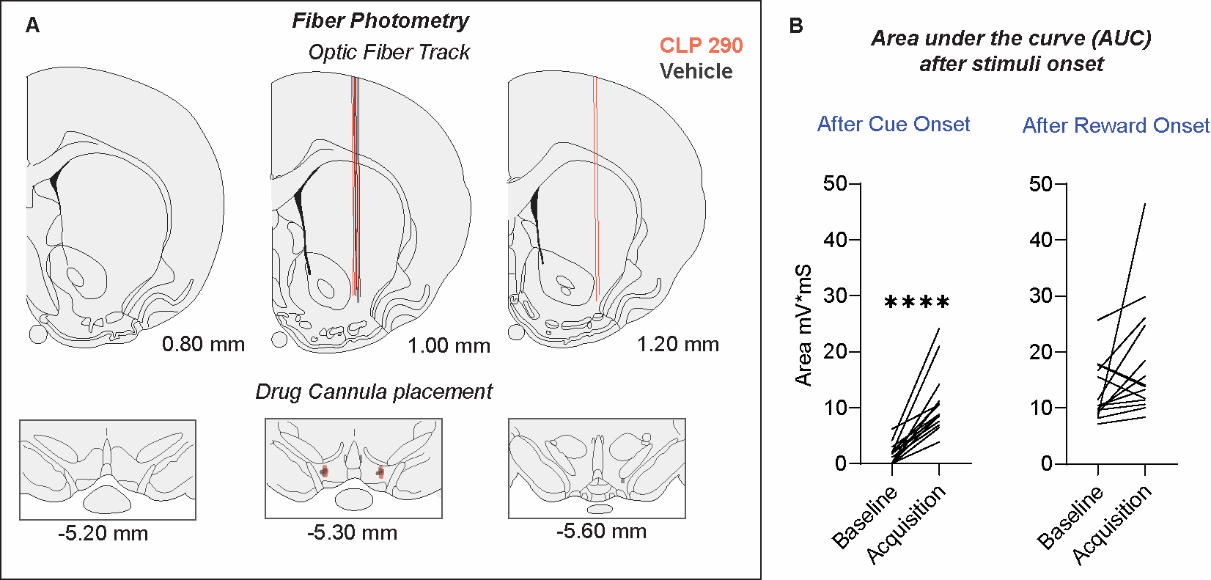

**Figure S6. Fiber photometry anatomical placements and fluorescence changes. Related to Figure 5.**

A) Anatomical placements of the optic fiber and drug delivery cannula for photometry were determined as described in the methods.

B) Cue-evoked GRAB_DA2M_ fluorescence significantly increased during acquisition compared to baseline of learning. In contrast, reward-evoked GRAB_DA2M_ fluorescence was not significantly different between baseline and acquisition of learning (p = 0.08, n = 14, paired t-test). ****p < 0.0001, paired t-test, n = 14.

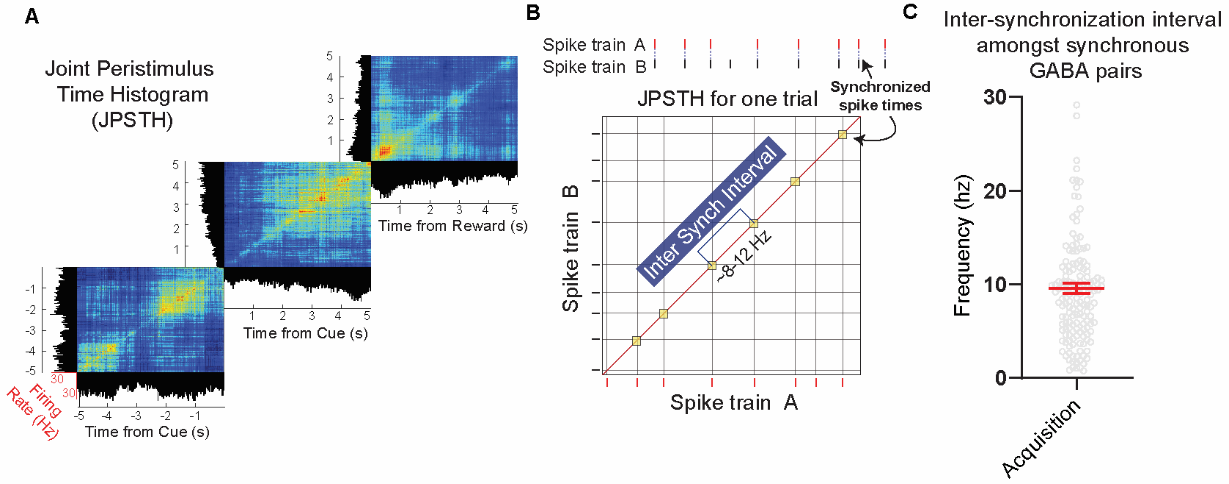

**Figure S7. VTA GABA neurons display millisecond time frame synchrony at around 10 hz. Related to Figure 6.**

A) Sample mean joint peristimulus time histograms (JPSTH) for pre-CS, CS, and US showing the correlation of firing activity in synchronized VTA GABA neurons over time.

B) Inter-synchronized intervals were calculated from JPSTHs as the frequency of coincident events during each trial (see methods).

C) During acquisition, inter-synchronized intervals between VTA GABA neurons were at approximately 10 Hz frequencies (Mean = 9.57 ± 0.55 Hz, n = 156 cells). Data are represented as mean ± SEM.

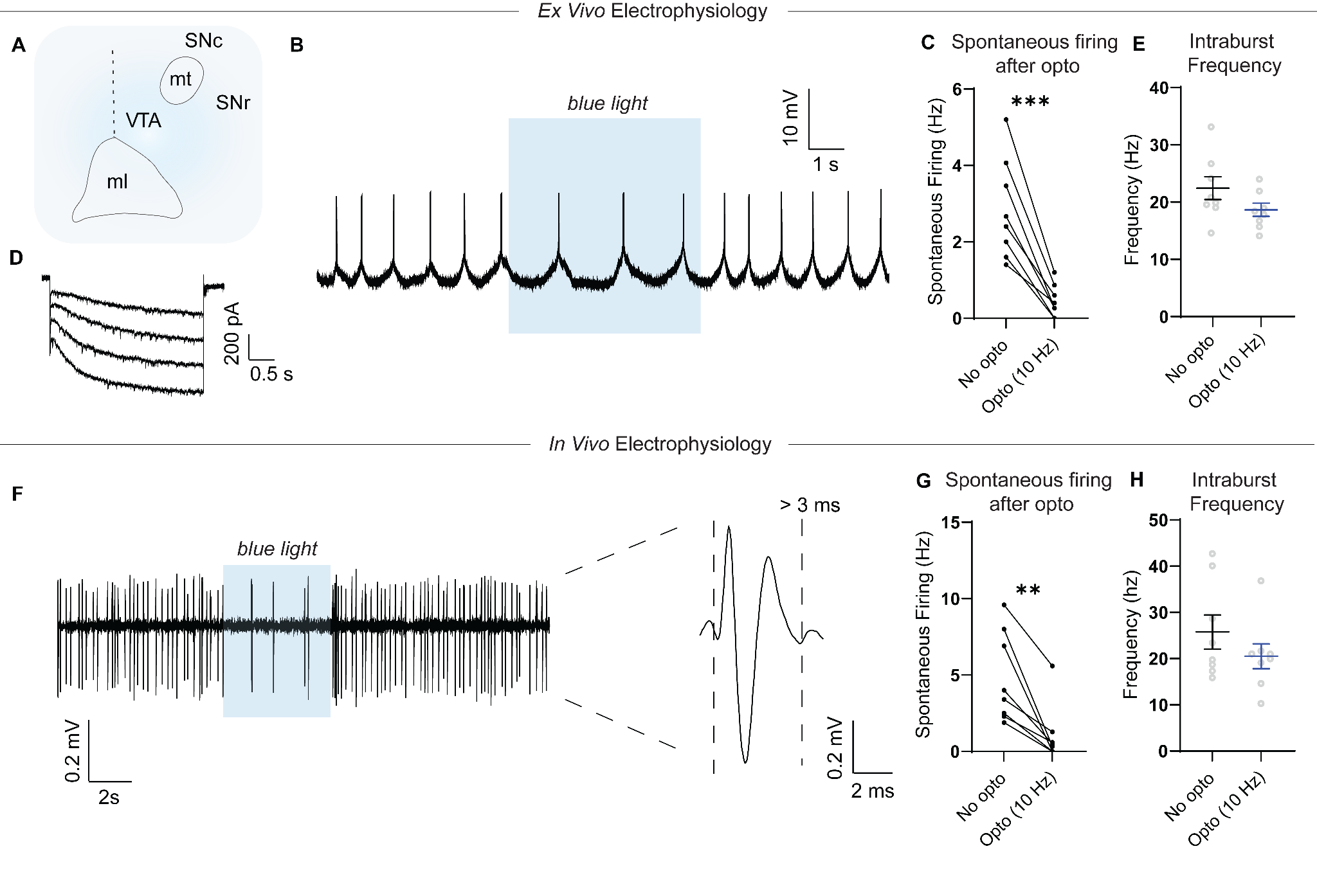

**Figure S8. DA neuron identification and intraburst frequency during *ex vivo* and *in vivo* electrophysiological recordings. Related to Figure 6.**

A) The lateral VTA in horizontal brain slices was identified as medial to the medial terminal nucleus of the accessory optic tract (mt) and lateral and rostral to the crest of the medial lemniscus (ml). Abbreviations: VTA = ventral tegmental area, SNc = substantia nigra pars compacta, SNr = substantia nigra pars reticulata, ml = medial lemniscus, mt = medial terminal nucleus of the accessory optic tract.
B) A representative recording from a DA neuron showing suppressed spontaneous firing activity after blue light stimulation of neighboring VTA GABA neurons. These DA neurons exhibited slow, spontaneous pacemaker firing characteristics (< 6 Hz).
C) Individual DA neuron responses to 10 Hz blue light application. Optogenetic stimulation of neighboring GABA responses significantly suppressed spontaneous DA neuron activity. ***p<0.005, paired t-test, n = 8 cells, 3 rats.
D) VTA DA neurons possessed large I_h_ (> 150 pA) current. I_h_ was recorded in whole-cell configuration using voltage step protocol (-40 mV to -110mV in 10 mV steps, 3 s duration, traces displayed for steps from -80 to -110). All calculations of the peak I_h_ amplitude were taken from a -60 mV to -110 mV hyperpolarizing step.
E) Intraburst frequencies during glutamate-induced burst firing showed no significant difference before and during 10 Hz light stimulation (p = 0.337, paired t-test, n = 8 cells).
F) Left: *In vivo* single-unit recordings in anesthetized rats showing suppressed firing activity of putative VTA DA neurons during blue light stimulation. Example traces for putative dopamine neurons showing low frequency spontaneous firing (<10 Hz) and broad triphasic action potential (>3 ms).
G) Putative DA neurons exhibited significantly suppressed spontaneous activity during 10 Hz blue light application **p < 0.01, paired t-test, n = 8 cells, 5 rats).
H) Intraburst frequencies during PPTg-induced burst firing showed no significant difference before and during 10 Hz light stimulation (p = 0.24, paired t-test, n = 8 cells).
